# GWAS of epigenetic ageing rates in blood reveals a critical role for *TERT*

**DOI:** 10.1101/157776

**Authors:** Ake T. Lu, Luting Xue, Elias L. Salfati, Brian H. Chen, Luigi Ferrucci, Daniel Levy, Roby Joehanes, Joanne M Murabito, Douglas P. Kiel, Pei-Chien Tsai, Idil Yet, Jordana T. Bell, Massimo Mangino, Toshiko Tanaka, Allan F. McRae, Riccardo E. Marioni, Peter M. Visscher, Naomi R. Wray, Ian J. Deary, Morgan E. Levine, Austin Quach, Themistocles Assimes, Philip S. Tsao, Devin Absher, James D. Stewart, Yun Li, Alex P. Reiner, Lifang Hou, Andrea A. Baccarelli, Eric A. Whitsel, Abraham Aviv, Alexia Cardona, Felix R. Day, John R.B. Perry, Ken K. Ong, Kenneth Raj, Kathryn L. Lunetta, Steve Horvath

## Abstract

DNA methylation age is an accurate biomarker of chronological age and predicts lifespan, but its underlying molecular mechanisms are unknown. In this genome-wide association study of 9,907 individuals, we found gene variants mapping to five loci associated with intrinsic epigenetic age acceleration (IEAA) and gene variants in 3 loci associated extrinsic epigenetic age acceleration (EEAA). Mendelian randomization analysis suggested causal influences of menarche and menopause on IEAA and lipid levels on IEAA and EEAA. Variants associated with longer leukocyte telomere length (LTL) in the telomerase reverse transcriptase gene (*TERT*) locus at 5p15.33 confer higher IEAA (P<2.7×10^-11^). Causal modelling indicates *TERT*-specific and independent effects on LTL and IEAA. Experimental hTERT expression in primary human fibroblasts engenders a linear increase in DNA methylation age with cell population doubling number. Together, these findings indicate a critical role for hTERT in regulating the DNA methylation clock, in addition to its established role of compensating for cell replication-dependent telomere shortening.

## INTRODUCTION

DNA methylation (DNAm) profiles of sets of cytosine phosphate guanines (CpGs) allow one to develop accurate estimators of chronological age which are referred to as “DNAm age”, “epigenetic age”, or the “epigenetic clock”. Across the life course the correlation between DNAm age and chronological age is greater than 0.95 ^1,2^. Individuals whose leukocyte DNAm age is older than their chronological age (“epigenetic age acceleration”) display a higher risk of all-cause mortality after accounting for known risk factors ^3-6^, and offspring of centenarians exhibit a younger DNAm age ^7^. Taken together, these findings suggest that DNAm age is a biomarker of biological age – a premise supported by associations of epigenetic age acceleration with cognitive impairment, neuro-pathology in the elderly ^8,9^, Down syndrome ^10^, Parkinson’s disease ^11^, obesity ^12^, HIV infection ^13^, and frailty ^14^, and menopause ^15^. However, DNAm age shows no apparent correlation with telomere length, whose pace of shortening in cultured somatic cells has been referred to as the ‘mitotic clock’. *In vivo,* DNAm age and telomere length appear to be independent predictors of mortality ^16^.

Here, we examine two widely used measures of epigenetic age acceleration: (a) intrinsic epigenetic age acceleration (IEAA), based on 353 CpGs described by Horvath (2013) ^2^, which is independent of age-related changes in blood cell composition, and (b) extrinsic epigenetic age acceleration (EEAA), an enhanced version of that based on 71 CpGs described by Hannum (2013) which up-weights the contribution of blood cell count measures ^1,6^. IEAA and EEAA are only moderately correlated (r=0.37) ^17^. IEAA measures cell-intrinsic methylation changes, exhibits greater consistency across different tissues, appears unrelated to lifestyle factors and probably indicates a fundamental cell ageing process that is largely conserved across cell types ^2,6^. By contrast, EEAA captures age-related changes in leukocyte composition and correlates with lifestyle^17^, yielding a stronger predictor of all-cause mortality ^6^. To understand mechanisms that explain DNAm age, we performed genome-wide association studies (GWAS) of IEAA and EEAA based on leukocyte DNA samples from almost ten thousand individuals.

## RESULTS

### GWAS meta-analyses for IEAA and EEAA

Genomic analyses were performed in as many as 9,907 individuals (aged 10-98 years), from 15 data sets, adjusted for chronological age and sex (**Supplementary Table 1, Fig. 1 and Supplementary note 1**). Eleven data sets comprised individuals of European ancestry (84.7%) and four comprised individuals of African (10.3%) or Hispanic ancestry (5.0%). GWAS genotypes were imputed to ~7.4 million variants using the 1000 genomes reference panel. Heritability estimates based on family relationships in one European ancestry cohort were 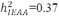 and 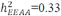, which are consistent with previous estimates in twins ^2^ and with IEAAEEAA those obtained in other tissues (e.g., adipose and brain) ^12,18,19^. SNP-based estimates of narrow sense heritability in our European ancestry cohorts were lower, 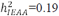 and 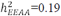 (**Supplementary Table 2**).

**Figure 1.**
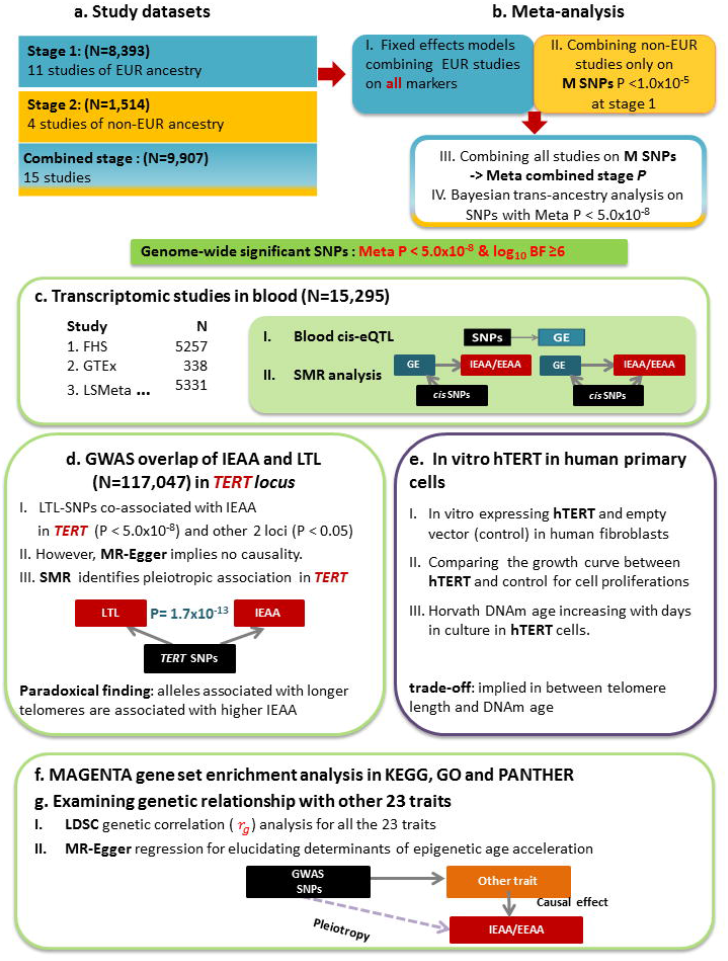
Roadmap of studying genetic variants associated with epigenetic age acceleration in blood Legend for Figure 1. The roadmap depicts our analytical procedures. Panel (a) describes the study sets were divided into two stages according to European (EUR) and non-European ancestry. Panel (b) indicates that stage 1 yielded GWAS summary data on all QC SNPs and the combined stage yielded GWAS summary data on the SNPs with Meta EUR P < 1.0×10^-5^ at stage 1. Genome-wide significant loci were determined based on the association results from the combined stage. Panel (c) describes our transcriptomic studies: (I) blood cis-eQTL to identify potential functional genes, (II) summary statistics based Mendelian randomization (SMR) to assess the causal associations between expression levels and IEAA (or EEAA). Panel (d) describes our detailed analysis in the *TERT* locus, which was implicated by our GWAS of IEAA. Bi-directional Mendelian randomization via MR-Egger analysis did not reveal a direct causal effect between leukocyte telomere length and IEAA. Our in vitro studies validate our genetic findings by demonstrating that *hTERT* over-expression promotes epigenetic ageing in Panel (e). To explore molecular pathways underlying epigenetic age acceleration, we conducted gene set enrichment analysis, as listed in Panel (f). Lastly, we performed LDSC genetic correlation between IEAA or EEAA and a broad category of complex traits, followed by MR-Egger regression analysis, as depicted in Panel (g). Abbreviation: GE= gene expression, LTL=leukocyte telomere length.

We first performed GWAS meta-analysis of IEAA and EEAA only in our European ancestry cohorts (N= 8,393). Variants with suggestive associations (P < 1.0×10^-5^) were then evaluated in non-European ancestry cohorts (N= 1,514), followed by a combined meta-analysis across the two strata (**Fig. 1a&b**). We found no evidence for genomic inflation in individual studies (λ*_GC_* = 0.99~1.06, **Supplementary Tables 3&4**) or in the European ancestry meta-analysis (λ*_GC_* = 1.03 ~ 1.05; LD score regression intercept terms β_0,*IEAA*_ = 1.004 and β_0,*EEAA*_ = 1.004; **Supplementary Fig. 1** and **Supplementary Table 2**). Variant associations were also adjusted for trans-ethnic heterogeneity using MANTRA software ^20^ which calculates a Bayes Factor (BF). Variants that met two criteria: P<5.0×10^-8^ and *log*_10_ *BF* ≥. 6 (approximately equivalent to P<5×10^-8^) were regarded as significant.

For IEAA, we identified 264 associated variants, mapping to five genomic loci (3q25.33, 5p15.33, 6p22.3, 6p22.2 and 17q22, **Table 1, Supplementary Table 5, Fig. 2 and Supplementary Fig. 2**). Conditional GCTA analyses revealed a secondary signal for IEAA at 6p22.3 (**Table 1, Supplementary Figs. 3c & 3h**). For EEAA, we identified 440 associated variants, mapping to three loci (4p16.3, 10p11.1 and 10p11.21; **Table 1**, **Supplementary Table 5**, **Fig. 2** and **Supplementary Fig. 4**), however the two lead SNPs, rs71007656 and rs1005277 at 10p11.1 and 10p11.21, respectively, are moderately correlated (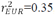, **Table 1**) and in a conditional model rs1005277, but not rs71007656, was associated with EEAA (**Supplementary Fig. 5b-c & e-f**). Associations were consistent across studies (**Supplementary Figs. 6 & 7**), except for at one locus (6p22.3: Cochran’s *I*^2^ =58%, MANTRA posterior probability of heterogeneity=0.64, **Table 1**). At the associated loci, each allele conferred between 0.41 to 1.68 years higher IEAA, or 0.59 to 0.74 years higher EEAA (**Table 1**). Analysis of published chromatin state marks ^21^ showed that most lead variants are in chromosomal regions that are transcribed in multiple cell lines (**Supplementary Fig. 8**). Two loci, 6p22.2 and 6p22.3, co-locate (within 1 Mb) with CpGs that contribute to the Horvath estimate of DNAm age (**Table 1** and **Supplementary Table 5**), and it is possible that these genotypic associations with IEAA arise from direct SNP effects on local methylation (**Supplementary note 2** and **Supplementary Figures 9 & 10**).

**Figure 2.**
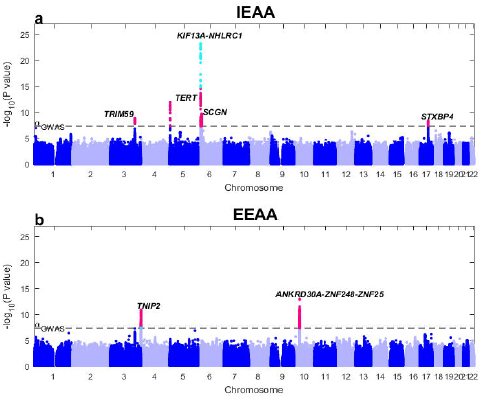
Genome-wide meta-analysis for intrinsic and extrinsic age acceleration in blood Legend for Figure 2. Manhattan plots for the meta-analysis p-values resulting from 15 studies comprised of 9,907 individuals. The y-axis reports log transformed p-values for (a) intrinsic epigenetic age acceleration (IEAA) or (b) extrinsic epigenetic age acceleration (EEAA). The horizontal dashed line corresponds to the threshold of genome-wide significance (*P*=5.0×10^-8^). Genome-wide significant common SNPs (MAF ≥ 5%) and low frequency SNPs (2% ≤ MAF < 5%) are colored red and cyan, respectively.

**Table 1:**
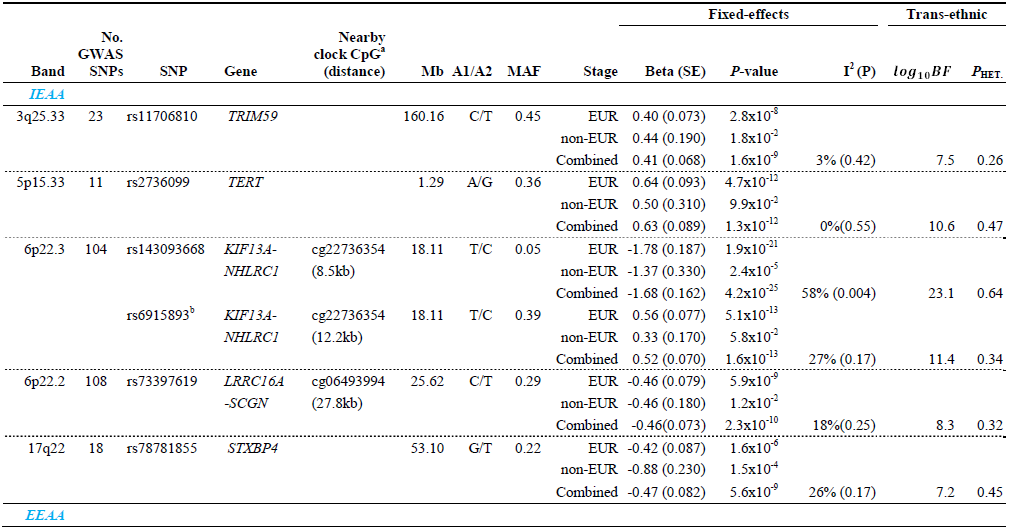
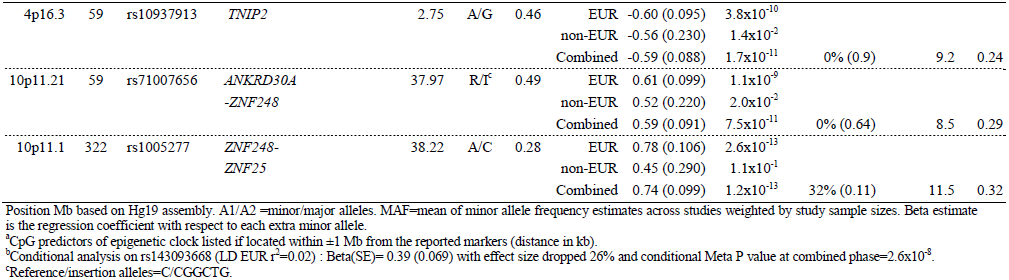
Meta-analysis of GWAS of epigenetic age acceleration in blood. **Legend** Lead SNPs at genome-wide significant (P<5.0x10^-8^) loci for IEAA or EEAA. Fixed effects meta-analysis was used to estimate the effect size (Beta) and standard error (SE) on IEAA or EEAA per minor allele. Trans-ethnic analyses using MANTRA^20^ present ethnicity-adjusted associations (log_10_ Bayes’ Factor (BF) and probability of heterogeneity across studies (*P*_HET._).

### Transcriptomic studies in leukocytes

To learn about potential functional consequences of these associations, we conducted cis-eQTL analysis for each locus associated with IEAA or EEAA, using data on leukocyte mRNA expression in up to 15,295 samples from five studies (**Fig. 1c; Methods)**. We identified 11 putative cis-eQTLs located in seven of the eight associated loci (**Supplementary Tables 6 & 7**). Each putative cis-eQTL was then analyzed by summary data-based Mendelian randomization (SMR), which infers the association between gene transcript levels and the outcome trait ^22^ (**Methods**). Three transcripts were associated with IEAA: *KPNA4* at 3q25.33, *TPMT* at 6p22.3 and *STXBP4* at 17q22; and three transcripts were associated with EEAA: *RNF4* at 4p16.3, and *ZNF25* and *HSD17B7P2* at 10p11.1 (**Table 2** and **Supplementary Table 8)**. Notably, *STXBP4*, encoding the syntaxin binding protein, is a reported locus for age at menarche ^23^, and our lead SNP for IEAA was also associated with age at menarche (rs78781855, P=0002). Consistent with our genetic analyses, blood transcript levels of several *cis*-acting genes correlate directly with chronological age, for example: *STXBP4*: r=-0.13, p=3×10^-9^; *RNF4*: r=-0.09, p=1.0×10^-5^; *ZNF25*: r=0.06, p=6.4×10^-3^ (**Supplementary Table 9**). Additional details can be found in **Supplementary note 3**.

**Table 2:** Summary of transcriptomic studies for loci associated with epigenetic age acceleration. **Legend** The table presents a total of six cis genes highlighted from transcriptomic study using three large-scale databases (N=10,906) including (1) FHS (N=5,257), (2) GTEx (N=338), and (3) LSMeta (5,311). Each cis gene exhibited significant cis-eQTL with several nearby GWAS SNPs at FDR q < 0.05 in at least one study and also showed a significant pleiotropic association in SMR analysis at P < 0.05 after Bonferroni correction. For each gene, we list the unadjusted P values (sign) from Meta GWAS for IEAA (or EEAA), cis-eQTL and SMR analysis. The column (sign) indicates the sign of Z statistic at each test while the test alleles were converted to the same alleles (with minor variants) for both Meta GWAS and cis-eQTL tests. The summary statistics of Meta GWAS and cis-eQTL are both based on the leading marker with the most significant P value in a given locus, according to the GWAS results.

| Band | SNP | Meta GWAS |  | cis-eQTL |  |  |  | SMR |
| --- | --- | --- | --- | --- | --- | --- | --- | --- |
|  |  | *P* (sign) |  | Database | Gene | *P* (sign) |  | *P* (sign) |
| IEAA |  |  |  |  |  |  |  |  |
| 3q25.33 | rs11706810 | 1.6x10<sup>-9</sup> | (+) | FHS | KPNA4 | 4.2x10<sup>-23</sup> | (+) | 5.1x10<sup>-5</sup> (+) |
| 6p22.3 | rs6915893 | 1.6x10<sup>-13</sup> | (+) | FHS | TPMT | 6.0x10<sup>-14</sup> | (+) | 1.1x10<sup>-7</sup> (+) |
|  |  |  | (+) | GTE<sub>x</sub> |  | 1.5x10<sup>-4*</sup> | (+) | 2.2x10<sup>-6</sup> (+) |
|  |  |  | (+) | LSMeta |  | 2.3x10<sup>-27</sup> | (+) | 3.5x10<sup>-7</sup> (+) |
| 17q22 | rs78781855 | 5.6x10<sup>-9</sup> | (-) | FHS | STXBP4 | 1.0x10<sup>-88</sup> | (-) | 6.2x10<sup>-4</sup> (+) |
| EEAA |  |  |  |  |  |  |  |  |
| 4p16.3 | rs2341303<sup>a</sup> | 6.5x10<sup>-11</sup> | (-) | LSMeta | RNF4 | 1.6x10<sup>-10</sup> | (-) | 2.0x10<sup>-3</sup> (+) |
| 10p11.21 | rs71007656 | 7.5x10<sup>-11</sup> | (+) | FHS | ZNF25 | 7.9x10<sup>-7</sup> | (+) | 2.8x10<sup>-3</sup> (+) |
|  |  |  |  | GTE<sub>x</sub> | HSD17B7P2 | 3.0x10<sup>-8</sup> | (+) | 6.9x10<sup>-6</sup> (+)<sup>b</sup> |
| 10p11.1 | rs1005277 | 1.2x10<sup>-13</sup> | (+) | GTE<sub>x</sub> | HSD17B7P2 | 1.1x10<sup>-5</sup> | (+) | 6.9x10<sup>-6</sup> (+)<sup>b</sup> |

Bands corresponding the position of Meta GWAS SNP.
\*FDR > 0.05 but FDR < 0.05 associated with other GWAS SNPs (**Supplementary Table 6**).
<sup>a</sup>The SNP rs2341303 is a surrogate of the leading marker rs10937913 in 4p16.3 (LD $r_{EUR}^2 = .98$ ).
<sup>b</sup>The SMR results are derived from the same model as the analysis used the cis SNPs of the gene *HSD17B7P2*.

### Alleles associated with higher IEAA at the *TERT* locus were also associated with longer telomere length

The *TERT* locus (in 5p15.33) harbored 11 genome-wide significant SNPs for IEAA but conditional analysis did not reveal any secondary signal (**Table 1**, **Fig. 3a**, and **Supplemental Table 5**). The leading SNP, rs2736099, was located in a region transcribed in human embryonic stem cells, induced pluripotent stem cells, and hematopoietic stem cells (**Supplementary Fig. 8b**), and each minor allele conferred 0.6 years higher IEAA (P=1.3×10^-12^; **Table 1, Supplementary Fig. 6b**). Our IEAA locus at *TERT* closely overlaps the reported GWAS locus for leukocyte telomere length (LTL ^24-26^, **Fig. 3b**). SMR analysis indicated that the association signals for LTL and IEAA at this *TERT* locus share the same underlying causal variant (as indicated by a non-significant HEIDI test, **Supplementary Table 10, Supplementary Fig. 11c**). Intriguingly, *TERT* alleles associated with a longer LTL (indicative of younger biological age) were robustly associated with increased IEAA (indicative of older biological age) (P ~ 1.0x10^-11^, **Table 3, Supplementary Table 11** and **Supplementary Figure 12**).

**Figure 3.**
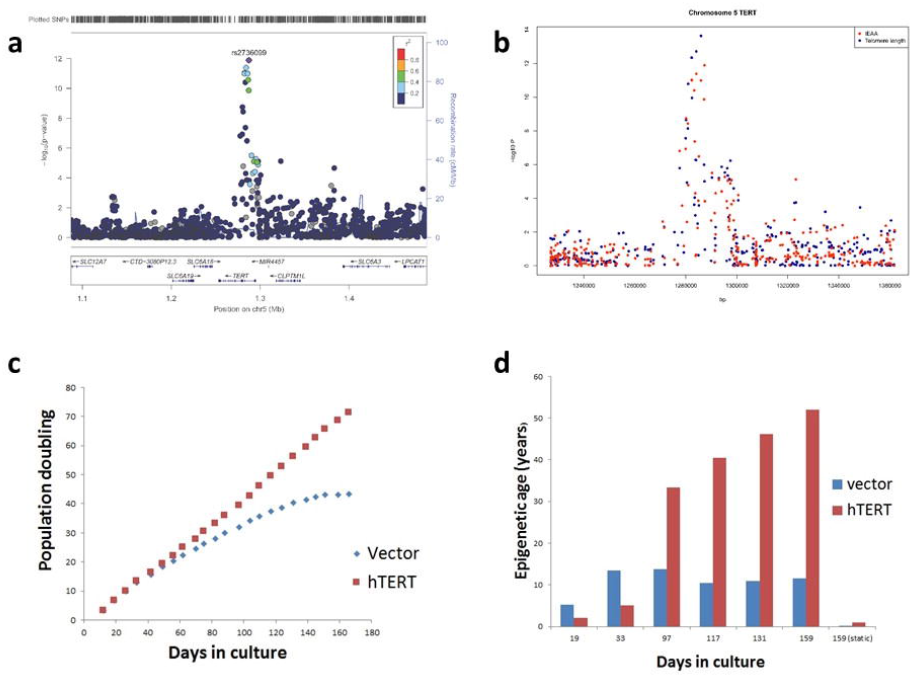
Genetic analysis of the 5p15.33 *TERT* locus and in vitro studies of hTERT in fibroblasts Legend for Figure 3. (a) Regional association plot of locus associated with IEAA. The y-axis depicts log-transformed meta-analysis P values across all studies 1-15. The colors visualize linkage disequilibrium (LD) r^2^ between rs2736099 (colored in purple) and neighboring SNPs. (b) *TERT*-locus association with IEAA (marked in red) overlaid with the association with telomere length given by Bojesen et al^24^ (marked in blue). Note that several SNPs in the TERT-locus are associated with both IEAA and leukocyte telomere length at a genome-wide significant level. (c) Growth of human primary fibroblasts represented as population doublings (y-axis) versus days in culture. (d) Adjusted epigenetic age of individual samples versus days in culture. The adjusted age estimate was defined as difference between DNAm age (Horvath method) minus 28 years, since the former exhibited a substantial offset in fibroblasts.

**Table 3:** Several leukocyte telomere length associated SNPs are also associated with intrinsic epigenetic age acceleration in blood and brain. **Legend** The table relates genome-wide significant association results of leukocyte telomere length (LTL) to three epigenetic age acceleration measures, IEAA in blood, IEAA in prefrontal cortex (PFCTX), and EEAA in blood. We queried the results of 14 SNPs across 10 distinct susceptibility loci associated with LTL from three large-scale studies: (I) meta-analysis association of LTL in chromosome 5 *hTERT* only (N=53,724)^24^, (II) a genome-wide meta-analysis of LTL (N=37,684)^25^, and (III) a genome-wide meta-analysis of LTL (N=26,089)^26^. Each row presents a genome-wide significant locus associated with LTL in a given study, except chromosome 16 *MPHOSPH6* and chromosome 20 *BCL2L1* just slightly below genome-wide significance and highlighted by the corresponding studies as major findings. The listed markers are the leading SNPs with the most significant P values associated with LTL at a given study and locus, sorted by chromosome and position. Effect sizes of LTL association refer to the change in telomere lengths (ΔTL). Telomere lengths were measured based on the relative telomere to single copy gene (T/S) ratios using standard qualitative PCR methods. A wide range of effect sizes for ΔTL across studies was due to different scaling approaches applied to the measurements. The effect sizes for each age acceleration measure are in units of year.

| Study | Chr | Gene | SNP | A1 | MAF | Effect size ($\beta$) | | | | Meta P value | | | |
| --- | --- | --- | --- | --- | --- | --- | --- | --- | --- | --- | --- | --- | --- |
|  |  |  |  |  |  | LTL<br>blood | IEAA<br>blood | IEAA<br>PFCTX | EEAA<br>blood | LTL<br>blood | IEAA<br>blood | IEAA<br>PFCTX | EEAA<br>blood |
| II | 2 | *ACYP2* | rs11125529 | A | 0.14 | 0.06 | -0.06 | 0.47 | -0.15 | $4.5 \times 10^{-8}$ | 0.5 | $9.9 \times 10^{-2}$ | $2.5 \times 10^{-1}$ |
| III | 3 | *PXK* | rs6772228 | A | 0.06 | -120 | 0.26 | -- | 0.20 | $3.9 \times 10^{-10}$ | $1.3 \times 10^{-1}$ | -- | $3.7 \times 10^{-1}$ |
| II | 3 | *TERC* | rs10936599 | T | 0.24 | -0.10 | 0.04 | 0.22 | -0.07 | $2.5 \times 10^{-31}$ | 0.7 | $3.5 \times 10^{-1}$ | $4.7 \times 10^{-1}$ |
| III | 3 | | rs1317082 | G | 0.23 | -77 | 0.04 | 0.22 | -0.08 | $1.3 \times 10^{-19}$ | 0.7 | $3.5 \times 10^{-1}$ | $4.6 \times 10^{-1}$ |
| II | 4 | *NAF1* | rs7675998 | A | 0.22 | -0.07 | -0.07 | -0.27 | 0.04 | $4.4 \times 10^{-16}$ | $4.4 \times 10^{-1}$ | $2.5 \times 10^{-1}$ | 0.7 |
| III | 5 | *TERT* | rs7726159 | A | 0.33 | 73 | 0.67 | 0.53 | 0.27 | $4.7 \times 10^{-17}$ | **$9.5 \times 10^{-12}$** | **$1.0 \times 10^{-2}$** | $6.4 \times 10^{-2}$ |
| I | 5 | | rs7705526 | A | 0.33 | 0.51 | 0.61 | 0.50 | 0.18 | $2.3 \times 10^{-14}$ | **$1.0 \times 10^{-11}$** | **$1.7 \times 10^{-2}$** | $1.9 \times 10^{-1}$ |
| II | 5 | | rs2736100 | C | 0.50 | 0.08 | 0.49 | 0.33 | 0.19 | $4.4 \times 10^{-19}$ | **$2.7 \times 10^{-11}$** | $1.0 \times 10^{-1}$ | $8.2 \times 10^{-2}$ |
| III | 10 | *OBFC1* | rs2487999 | T | 0.14 | 100 | 0.22 | 0.40 | 0.36 | $4.2 \times 10^{-14}$ | **$3.7 \times 10^{-2}$** | $2.5 \times 10^{-1}$ | **$8.8 \times 10^{-3}$** |
| II | 10 | | rs9420907 | C | 0.20 | 0.07 | 0.26 | 0.34 | 0.20 | $6.9 \times 10^{-11}$ | **$4.1 \times 10^{-3}$** | $2.4 \times 10^{-1}$ | $9.9 \times 10^{-2}$ |
| II | 16 | *MPHOSPH6* | rs2967374 | A | 0.22 | 0.05 | 0.23 | 0.37 | 0.12 | $2.7 \times 10^{-7}$ | **$7.8 \times 10^{-3}$** | $1.2 \times 10^{-1}$ | $2.9 \times 10^{-1}$ |
| II | 19 | *ZNF208* | rs8105767 | G | 0.32 | 0.05 | -0.05 | 0.35 | -0.13 | $1.1 \times 10^{-9}$ | 0.5 | $1.1 \times 10^{-1}$ | $1.7 \times 10^{-1}$ |
| III | 20 | *BCL2L1* | rs6060627 | T | 0.34 | 36 | -0.13 | -0.14 | 0.11 | $6.5 \times 10^{-7}$ | $7.7 \times 10^{-2}$ | 0.5 | $2.6 \times 10^{-1}$ |
| II | 20 | *RTEL1* | rs755017 | G | 0.15 | 0.06 | 0.03 | -0.03 | -0.08 | $6.7 \times 10^{-9}$ | 0.8 | 0.9 | 0.5 |

--denote not available. P values associated with age acceleration marked in bold if $< 0.05$ and red if $< 5.0 \times 10^{-8}$ .
Chr=chromosome; A1: reference allele; MAF=minor allele frequency; Effect sizes corresponding additive models.

Other known GWAS LTL signals (at 10q24.33 near *OBFC1* and at 16q23.3 near *MPHOSPH6*), also exhibited modest associations with IEAA (4.1×10^-3^ ≤ P ≤ 3.7×10^-2^), but others such as the gene *TERC* (on 3q26.2) encoding the telomerase RNA component, showed no association (**Table 3**). Using the lead variants for each trait in MR-Egger analyses^27^, we found pleiotropic effects specific to *TERT* on LTL and IEAA without evidence for a causal relation between LTL and IEAA (P =0.7, **Table 4** and **Supplementary Table 12**).

IEAA-associated SNPs in the *TERT* locus also exhibit modest associations with epigenetic age acceleration in the prefrontal cortex (P < 0.05, **Table 3**). We found no association between LTL-related SNPs and epigenetic age acceleration in other brain regions, which were represented by fewer brain samples (**Supplementary Table 13**).

### hTERT is required for DNAm ageing in human primary cells

As we were unable to functionally link IEAA to *TERT* through *cis* eQTL analysis at 5p15.33, we examined the effects of experimental-induced hTERT expression on IEAA in a primary human cell culture model. We introduced a *TERT-*expressing vector or empty vector (as control) into primary fibroblasts isolated from human neonatal foreskin. Transduced *TERT*-expressing and non-*TERT* cells were cultured in parallel. After reaching confluence, the cells were harvested, counted, seeded into fresh plates, and profiled using the Illumina Infinium 450K DNA methylation array.

While non-*TERT* cells senesced after ~150 days, *TERT*-expressing cells continued to proliferate unabated at a constant rate with the time in culture **(Fig. 3c)**. Single time point analyses (**Fig. 3d**) showed that *TERT*-expressing cells exhibited a linear relationship between time in culture and the Horvath estimate of DNAm age (equivalent to a DNAm age of 50 years at 150 days), whereas in non-*TERT* cells DNAm age plateaued (equivalent to a DNAm age of 13 years) in spite of continued proliferation to the point of replicative senescence. Notably, DNAm age did not increase in *TERT*-expressing cells that received regular media change but were not passaged throughout the entire observation period of 170 days (right most bar in **Fig. 3d**). These cells were not senescent, given that their subsequent passaging resulted in normal proliferation. In multivariable regression analysis, the associations of DNAm age with cell passage number and cell population doubling number were highly modified by *TERT*-expression (P-interaction: P=1.6×10^-6^ and P=4.0×10^-5^, respectively; **Supplementary Table 14**). In the absence of *TERT*-expression, DNAm age did not increase with cell passage number, cell population doubling number, or time in culture.

### Other putative determinants of epigenetic age acceleration

To systematically elucidate possible further biological processes that influence epigenetic age acceleration, we tested our full genome-wide association statistics for IEAA and EEAA using a number of approaches. First, we used MAGENTA ^28^ (**Methods)** to identify biological pathways that are enriched for genes that harbor associated variants. For IEAA, nuclear transport (FDR=0.017), Fc epsilon RI signaling (FDR=0.027), and colorectal cancer processes (FDR=0.042) were implicated. For EEAA, mRNA elongation (FDR 0.011), mRNA transcription (FDR=0.018), and neurotrophin signaling pathway (FDR=0.042) were implicated (**Supplementary Table 15**).

Second, we explored the genetic correlations (*r_g_*) between IEAA or EEAA and several other phenotypes using LD score regression analysis of summary level GWAS data ^29^ (**Methods and Fig. 1g**). We observed moderate positive genetic correlation between IEAA and EEAA (*r_g_* =0.5, P*_rg_* =8.9×10^-3^). IEAA showed weak positive genetic correlations with central adiposity (waist gcircumference; waist-to-hip ratio) and metabolic disease-related traits, and EEAA showed stronger positive genetic correlations with central adiposity (*r_g_* =0.27 with waist-to-hip ratio, P=8.0×10^-8^) and metabolic disease-related traits (**Table 4** and **Supplementary Table 16**). IEAA and EEAA also showed modest inverse genetic correlations or trends with age at menopause. Third, we performed MAGENTA based hypergeometric analyses to test whether the top 2.5% and 10% of genes enriched for GWAS associations with IEAA or EEAA overlap with the top enriched genes for a range of complex traits (**Methods**). This analysis suggested several additional possible genetic overlaps, including Huntington disease onset^30^ and bipolar disorder with IEAA, schizophrenia with EEAA, and age-related dementia^18^ with both IEAA and EEAA (**Supplementary Table 17**).

**Table 4:** Genetic correlations with and causal effects of other complex traits on epigenetic age acceleration. **Legend** Results from cross trait LDSC genetic correlation and Mendelian randomization Egger regression (MR-Egger) analyses for IEAA and EEAA are presented. The traits are ordered by category (I) GWAS of anthropometric traits conducted by GIANT consortium, (II) GWAS of lipid, metabolic, and inflammatory outcomes and diseases, (III) GWAS of neurodegenerative and neuropsychiatric disorders, (IV) cognitive functioning and educational attainment traits, and (V) longevity, reproductive ageing and mitotic clock related traits. Complete results are presented in **Supplementary Tables 16, 18** and **19.** We list the sample size of a study trait, Genetic correlation (r_g_) and its P value (P_rg_) as well the estimate of causal effect (β _causal_) and its P value (P_causal_) from MRs-Egger regression. A 2-color scale (blue to red) applies to r_g_ in a range of [-1, 1], when the trait exhibits P_rg_ ~ ≤ 0.05.

|  |  | IEAA |  |  |  | EEAA |  |  |  |
| --- | --- | --- | --- | --- | --- | --- | --- | --- | --- |
|  |  | Genetic |  | MR-Egger |  | Genetic |  | MR-Egger |  |
|  |  | correlation |  | regression |  | correlation |  | regression |  |
| Trait | N | $r_g$ | $P_{r_g}$ | $\beta_{causal}$ | $P_{causal}$ | $r_g$ | $P_{r_g}$ | $\beta_{causal}$ | $P_{causal}$ |
| Category (I) |  |  |  |  |  |  |  |  |  |
| Waist circumference (cm) | 232,101 | 0.09 | $2.4 \times 10^{-2}$ | 1.99 | $9.6 \times 10^{-2}$ | 0.24 | $9.9 \times 10^{-7}$ | -1.48 | 0.3 |
| Waist-to-hip ratio | 212,243 | 0.13 | $1.0 \times 10^{-2}$ | -0.31 | 0.9 | 0.27 | $8.0 \times 10^{-8}$ | -3.36 | 0.3 |
| BMI (SD) | 339,224 | -0.01 | 0.9 | 1.40 | 0.056 | 0.08 | 0.3 | 0.40 | 0.7 |
| Height (cm) | 133,453 | -0.02 | 0.7 | 0.004 | $1.3 \times 10^{-1}$ | 0.13 | $2.8 \times 10^{-3}$ | 0.004 | $1.6 \times 10^{-1}$ |
| Category (II) |  |  |  |  |  |  |  |  |  |
| High density lipoprotein (SD) | 188,577 | -0.08 | $1.1 \times 10^{-2}$ | 0.37 | $1.2 \times 10^{-1}$ | -0.12 | $2.0 \times 10^{-2}$ | 0.11 | 0.7 |
| Triglyceride (SD) | 188,577 | 0.10 | $3.4 \times 10^{-2}$ | 0.61 | $3.0 \times 10^{-2}$ | 0.16 | $1.0 \times 10^{-4}$ | 0.37 | $8.0 \times 10^{-2}$ |
| Type 2 diabetes | 69,033 | 0.16 | $3.5 \times 10^{-2}$ | 0.08 | 0.8 | 0.09 | $1.8 \times 10^{-1}$ | 0.07 | 0.8 |
| Inflammatory bowel disease | 34,652 | 0.12 | $2.5 \times 10^{-2}$ | 0.13 | 0.5 | 0.08 | $1.2 \times 10^{-1}$ | -0.35 | $1.5 \times 10^{-1}$ |
| Crohn's disease | 20,883 | 0.12 | $4.7 \times 10^{-2}$ | 0.10 | 0.4 | 0.10 | $6.9 \times 10^{-2}$ | -0.29 | $7.6 \times 10^{-2}$ |
| Category (III & IV) |  |  |  |  |  |  |  |  |  |
| AMD subtype | 45,818 | 0.03 | 0.2 | -0.05 | 0.6 | 0.05 | $2.1 \times 10^{-2}$ | -0.04 | 0.7 |
| Educational attainment (years) | 328,917 | -0.01 | 0.8 | 2.35 | 0.3 | -0.13 | $1.0 \times 10^{-3}$ | -4.63 | $7.8 \times 10^{-2}$ |
| Category (V) |  |  |  |  |  |  |  |  |  |
| Age at menarche (years) | 252,514 | -0.02 | 0.8 | -1.03 | $4.1 \times 10^{-3}$ | -0.03 | 0.7 | -0.17 | 0.7 |
| Age at menopause (years) | 69,360 | -0.12 | $5.4 \times 10^{-2}$ | -0.43 | $3.5 \times 10^{-3}$ | -0.17 | $2.0 \times 10^{-3}$ | -0.21 | 0.3 |
| Leukocyte telomere length (T/S) | 37,684 | 0.18 | $9.7 \times 10^{-2}$ | 1.80 | 0.7 | -0.16 | $1.4 \times 10^{-1}$ | 3.20 | 0.3 |

Finally, all the study traits, were tested using MR-Egger which, by modelling the reported top genetic signals for each candidate trait, estimates the likely causal influence of that trait on IEAA or EEAA ^27^ (**Methods**). Nominally significant causal relationships on higher IEAA and EEAA were found for low density lipoprotein (LDL) and total cholesterol levels (P<0.05, **Supplementary Tables 18 & 19**) and for triglyceride levels on IEAA (P=3.0×10^-2^, **Table 4**). Earlier menarche and menopause were associated with higher IEAA; each 1-year earlier age at menarche was associated +1.03 years higher IEAA (P=4.1×10^-3^) and each 1-year earlier age at menopause was associated +0.43 years higher IEAA (P=3.5×10^-3^) (**Table 4**).

## DISCUSSION

This large genomic study provides several mechanistic insights into the regulation of epigenetic ageing, including apparently opposing roles for *TERT* on DNAm age and LTL. *TERT* encodes the catalytic subunit of telomerase, which counters telomere shortening during cell division ^31^. *TERT* also possesses activities unrelated to telomere maintenance, such as in DNA repair, cell survival, protection from apoptosis/necrosis, stimulation of growth ^32-34^ and cell proliferation, possibly by decreasing p21 production ^35^. Here, we show an additional pleiotropic role of *TERT* on advancing cell intrinsic DNAm age during cell proliferation. Our findings provide an explanation for the previously reported rapid rate of DNAm ageing during embryonic development and early postnatal life, which are stages of rapid organismal growth accompanied by high levels of *TERT* activity and cell division ^36^ ^37,38^.

Our paradoxical finding that *TERT* alleles associated with longer telomeres are associated with higher IEAA is substantiated by our *in vitro* evidence that *TERT* expression promotes DNAm age; this might suggest a potential trade-off between telomere length and epigenomic maintenance systems. However, we found no evidence for a broader causal inter-relationship between telomere length and IEAA, consistent with the lack of phenotypic association between these traits in our studies (WHI: r = -0.05, p=0.16; FHS: r=0.0, p=0.99, ref ^39^) and in previous reports ^14,16^. Furthermore, while critically short telomere length is a well-established trigger of replicative senescence ^36,40-42^, the functional consequences of epigenetic ageing are yet unknown. Our experimental data suggest that epigenetic ageing is not a determinant or marker of cell replicative senescence, since *TERT*-expressing cells continued to proliferate unabatedly despite well-advanced DNAm age, and non-*TERT*-expressing cells exhibited no DNAm age increase even at days 120-170 when proliferation had ceased. Rather, *TERT* suppression appears to allow cells to record their proliferation history during stages of development.

Large-scale cross-sectional cohort studies have previously reported associations between epigenetic age acceleration (IEAA and EEAA) and body mass index and measures of insulin resistance ^17^. Our genetic correlation analyses (between the measures of age acceleration and complex phenotypes) indicate that some of these associations may arise in part from shared genetic variants. Moreover, our Mendelian randomization analyses provided tentative evidence for causal influences for blood lipid levels, but not for adiposity, on IEAA and EEAA.

Finally, we found evidence for causal influence of earlier ages at menarche and menopause on higher IEAA. The directionally-concordant influences of menarche and menopause, which signal the onset and cessation, respectively, of reproductive capacity, together with the lack of influence of any measure of adiposity on IEAA, suggest an effect of some yet identified driver of reproductive ageing on DNAm ageing. These findings, which suggests that sex steroids affect epigenetic ageing, are consistent with previously reported associations regarding early menopause timing and higher IEAA in blood ^15^. While menopausal hormone therapy was not found to be associated with IEAA in blood, it was found to be associated with younger epigenetic age acceleration in buccal epithelium ^15^. The effect of menopause is consistent with reported anti-ageing effects of sex hormone therapy on buccal cells and the pro-ageing effect of the surgical ovariectomy in blood ^15^. Early age at menarche, a widely studied marker of the timing of puberty in females, is associated with higher risks for diverse diseases of ageing ^43^. Our findings indicate epigenetic ageing as a possible intrinsic mechanism that underlies the recently described link between menarche age-lowering alleles and shorter lifespan ^44^.

## ONLINE METHODS

### GWAS Cohorts

GWAS meta-analysis was performed on 9,907 individuals across 15 studies (**Supplementary Table 1)** coming from eight cohorts: Framingham Heart Study (FHS), TwinsUK, Women’s Health Initiate (WHI), European Prospective Investigation into Cancer–Norfolk (EPIC-Norfolk), Baltimore Longitudinal Study of Aging (BLSA), Invecchiare in Chianti, ageing in the Chianti Area Study (inCHIANTI), Brisbane Systems Genetics Study (BSGS), and Lothian Birth Cohorts of 1921 and 1936 (LBC) (**Supplementary note 1**). Eleven data sets comprised individuals of European ancestry (EUR, 84.7%) and four data sets comprised individuals of African ancestry (AFR, 10.3%) or Hispanic ancestry (AMR, 5.0%). Age range was 10-98 years (69% females).

### DNA methylation age and measures of age acceleration

By contrasting the DNAm age estimate with chronological age, we defined measures of epigenetic age acceleration that are uncorrelated with chronological age. We distinguished two types of measures of epigenetic age acceleration in blood: cell-intrinsic and extrinsic epigenetic measures, which are independent of, or enhanced by blood cell count information, respectively. Intrinsic epigenetic age acceleration (IEAA) is defined as the residual resulting from regressing the Horvath’s estimate of epigenetic age on chronological age and measures of blood cell counts. Extrinsic epigenetic age acceleration (EEAA) does depend on blood cell counts because it is defined by up-weighting the blood cell count contributions to the Hannum based measure of age acceleration. Thus, EEAA captures both age-related changes in blood cell types as well as cell-intrinsic age-related changes in DNAm levels.

IEAA and EEAA are based on the DNAm age estimates from Horvath ^2^ (353 CpG markers) and from Hannum ^1^ (71 CpGs), respectively. Mathematical details and software tutorials for estimating epigenetic age can be found in Horvath (2013) ^2^. To estimate “pure” epigenetic ageing effects that are not influenced by differences in blood cell counts (cell-intrinsic epigenetic age acceleration, IEAA), we obtained the residual resulting from a multivariate regression model of epigenetic age on chronological age and various blood immune cell counts (naive CD8+ T cells, exhausted CD8+ T cells, plasma B cells, CD4+ T cells, natural killer cells, monocytes, and granulocytes) imputed from methylation data. Our measure of EEAA is defined using the following three steps. First, we calculated the epigenetic age measure from Hannum et al ^1^, which already correlated with certain blood cell types ^3^. Second, we increased the contribution of immune blood cell types to the age estimate by forming a weighted average of Hannum’s estimate with 3 cell types that are known to change with age: naïve (CD45RA+CCR7+) cytotoxic T cells, exhausted (CD28-CD45RA-) cytotoxic T cells, and plasma blasts using the Klemera-Doubal approach ^45^. The weights used in the weighted average are determined by the correlation between the respective variable and chronological age ^45^. The weights were chosen on the basis of the WHI data. Thus, the same (static) weights were used for all data sets. Third, EEAA was defined as the residual variation resulting from a univariate model regressing the resulting age estimate on chronological age. By construction, EEAA is positively correlated with the estimated abundance of exhausted CD8+ T cells, plasma blast cells, and a negative correlated with naive CD8+ T cells. Blood cell counts were estimated based on DNA methylation data as described in the next section. By construction, the measures of EEAA track both age related changes in blood cell composition and intrinsic epigenetic changes. Both the intrinsic and extrinsic DNAm age measures were correlated highly with chronological age within each contributing cohort (0.63 ≤ *r* ≤ *0.*97, S**upplementary Table 1**), except for the two Lothian *birth* cohorts whose participants were born in one of two single years and hence had a small age range at testing. Conversely, by design, our measures of DNAm *age acceleration*, IEAA and EEAA are unrelated with chronological age.

The measures of epigenetic age acceleration are implemented in our freely available software (https://dnamage.genetics.ucla.edu) ^2^.

### Estimating blood cell counts based on DNA methylation levels

We estimated blood cell proportions using two different software tools. Houseman’s estimation method ^46^, which is based on DNA methylation signatures from purified leukocyte samples, was used to estimate the proportions of cytotoxic (CD8+) T cells, helper (CD4+) T, natural killer, B cells, and granulocytes. The software does not identify the type of granulocytes in blood (neutrophil, eosinophil, or basophil) but neutrophils tend to be the most abundant granulocyte (~60% of all blood cells compared with 0.5-2.5% for eosinophils and basophils). The advanced analysis option of our epigenetic age calculator software was used to estimate the percentage of exhausted CD8+ T cells (defined as CD28-CD45RA-) and the number (count) of naïve CD8+ T cells (defined as CD45RA+CCR7+) as described in Horvath et al.^13^ These DNAm based estimates of blood cell counts are highly correlated with corresponding flow cytometric measures 47

### GWAS meta-analysis

Our GWAS meta-analysis involved approximately 7.4 million SNPs/INDEL variants, which were genotyped and imputed markers with the 1000 genomes haplotype reference panel. Prior to imputation, SNP quality was assessed by estimating minor allele frequency (MAF), Hardy-Weinberg equilibrium (HWE), and missingness rates across individuals (**Supplementary Table 4)**. The individual studies used IMPUTE2 ^48,49^ with haplotypes phased using SHAPEIT ^50^ or MaCH ^51^ phased using Beagle ^52^ or Minimac ^48^ to impute SNP and INDEL markers based on the 1000 Genomes haplotypes released in 2011 June or 2012 March. The quality of imputed markers was assessed by the Info measure > 0.4 (in IMPUTE2) or R^2^ > 0.3 (in Minimac), and HWE P > 1.0x10^-6^. To increase resolution for SNP association, a few genomic regions in the FHS cohort were also imputed based the Haplotype Reference Consortium (N=64,976)^53^. FHS used linear mixed models to account for pedigree structure via a kinship matrix, as implemented in R “lmekin” package. The BSGS cohort used Merlin/QTDT ^54^ for family-based association analysis. For other association analyses, we regressed the age acceleration trait values on estimated genotype dosage (counts of test alleles) or (2) expected genotype dosage, implemented in Mach2QTL^55^, SNPTEST^56^, and PLINK. All association models were adjusted for sex, to account for the higher epigenetic age acceleration in men than women ^47^, and also for PCs as needed. We included variants with MAF ≥ 2%. SNPs were removed from an individual study if they exhibited extreme effects (absolute regression coefficient β >30, **Supplementary Table 4**).

We divided the meta-analysis into two since IEAA and EEAA differ across racial/ethnic groups ^47^. In one arm, we performed GWAS meta-analysis of IEAA and EEAA, focusing on individuals of European ancestry (N= 8,393, studies 1-11 in **Supplementary Table 1**). We required a marker present in at least 5 study data sets and combined the coefficient estimates β from each study using a fixed-effects meta-analysis model weighted by inverse variance, as implemented in the software Metal ^57^. In the other arm, each SNP with suggestive association (P < 1.0×10^-5^) in Europeans was subsequently evaluated in individuals of non-European ancestry (N= 1,514, studies 12-15 in **Supplementary Table 1**). A further meta-analysis combined the GWAS findings from the two ancestries. We removed SNPs from the meta-analysis if they exhibited highly significant heterogeneity across studies (Cochran Q I^2^ *P*-value ≤ 0.001), or (2) co-located with CpG from the DNAm age predictors according to the Illumina annotation file for the Illumina Infinium 450K array. We analyzed additional SNPs across all study sets to arrive at summary statistics at the combined stage, which were needed for our summary statistics based Mendelian randomization analyses. The quality of SNPs was also assessed using the Cochran Q I^2^ *P*-value.

### Linkage disequilibrium analysis

Regional SNP association results were visualized with the software LocusZoom^58^. All linkage disequilibrium (LD) estimates presented in this article were calculated using individuals of European ancestry from the 1000 genomes reference panel (released in Oct 2012).

### Conditional analysis based on GCTA

The conditional analysis of the GCTA software ^59-61^ was used to test whether a given genetic locus harbored multiple independent causal variants. We conditioned on the leading SNP with the most significant meta-analysis P value (**Table 1**). As reference panel for inferring the LD pattern we used the N=379 individuals with European ancestry from the 1000 genomes panel released in December 2013. We defined a SNP as having an additional association if it remained significant (P<5×10^-8^) after conditioning on the leading SNP and also met the additional criterion log_10_BF ≥ 6 for a significant trans-ethnic association.

### Chromatin state annotations

For each leading SNP/variant of a significant locus, we used the UCSC genome browser to display the primary chromatin states across 127 cell/tissue lines at 200bp resolution (**Supplementary Fig. 8**). The n=127 diverse cell/tissue lines were profiled by the NIH RoadMap Epigenomics ^21^ (n=111) and ENCODE projects ^62^ (n=16). We used a 15-state chromatin model (from ChromHMM) which is based on five histone modification marks ^21^.

### Annotations for genome-wide significant variants

We used the HaploReg (version 4.1) tool ^63^ to display characters of genome-wide significant variants including conserved regions by GERP^64^ and SiPhy^65^ scores, DNase tracks, involved proteins and motifs, GWAS hits listed in NHGRI/EBI and functional annotation listed in dbSNP database, as summarized in **Supplementary Table 5**.

### Leucocyte *cis*-eQTL analyses

To evaluate cis-eQTL in blood, our cis-eQTL study leveraged a large-scale blood expression data (n=15,295) that came from five broad categories of data. The first category involved a large-scale eQTL analysis in 5,257 individuals collected from the FHS pedigree cohort (of European ancestry)^66^. Linear mixed models were performed for the eQTL analysis, adjusted for family structure via random effects and adjusted gender, age, blood cell counts, PCs, and other potential confounders via fixed effects. The analysis was carried out using the *pedigreemm* package of R. The second category involved the significant *cis*-eQTL, released from GTEx (version 6 in 2015) ^67^. The expression data from GTEx involve multiple tissues from 449 individuals of mostly (>80%) European ancestry. We used the *cis* eQTL results evaluated in 338 blood samples. The downloaded *cis*-eQTL results only list significant results (FDR q < 0.05), according to a permutation test based threshold that corrected for multiple comparisons across genes and tissue types. The third category involved the *cis*-eQTL results from LSMeta^68^ which was a large-scale eQTL meta-analysis in 5,331 blood samples collected from 7 studies including our study cohort inChianti. We downloaded the *cis*-eQTL results from <u>http://genenetwork.nl/bloodeqtlbrowser/</u>.

The fourth and fifth categories were discovery and replication samples from an eQTL analysis in peripheral blood ^69^, respectively. The publicly released *cis*-eQTL results only involved the 9,640 most significant (FDR q < 0.01) cis-eQTL results (corresponding to 9,640 significant genes) from the discovery sample. The fourth category involved the expression data of 2,494 twins from the NTR cohort. The fifth category involved 1,895 unrelated individuals from the NESDA cohort.

For all five categories of blood data, the cis-window surrounding each SNP marker was defined as ±1 Mb. We defined a significant cis-eQTL relationship by imposing the following criteria: a) FDR q < 0.05 for categories 1 -3, b) FDR q < 0.01 for categories 4 & 5.

### SMR analysis

SMR^22^ uses SNPs as instrumental variables to test for a direct association between an exposure variable and an outcome variable, irrespective of potential confounders. Unlike conventional Mendelian randomization analysis, the SMR test uses summary-level data for both SNP-exposure and SNP-outcome that can come from different GWAS studies ^22^. We tested the expression levels of the eleven candidate genes identified in our leucocyte cis-eQTL analysis. SMR defines a pleiotropic association as association between gene expression and a test trait due to pleiotropy or causality (**Supplementary Fig**. **11a-b**). A significant SMR test p-value does not necessarily mean that gene expression and the trait are affected by the same underlying causal variant, as the association could possibly be due to the top associated cis-eQTL being in LD with two distinct causal variants. Zhu et al (2016) define the scenario of several causal variants, which is of less biological interest than pleiotropy, as “linkage” and proposed a statistical test “HEIDI” for distinguishing it from pleiotropy. The null hypothesis of the HEIDI test corresponds to desirable causal scenarios). Thus, a non-significant p value (defined here as P ≥ 0.01) of the HEIDI test is a desirable finding. Conversely, a significant HEIDI test p-value indicates that at least two linked causal variants affect both gene expression and epigenetic age acceleration (**Supplementary Fig. 11c**).

To test the association of a given gene expression with age acceleration, we used summary level *cis*-eQTL results from (1) FHS, (2) GTEx and (3) LSMeta (total: N=10,906).

We included the *cis* SNPs (with MAF ≥ 0.10) within a test gene (± 1Mb). We selected the *cis* SNPs as instrumental markers as follows: *cis*-eQTL FDR < .05 for GTEx, *cis*-eQTL P=1×10^-6^ for the two large-scale studies: LSMeta and FHS. All significant SNP-gene pairs were subjected to the HEIDI analysis. The analysis involves the summary data of the *cis* SNPs surrounding the instrumental markers and the LD pattern evaluated from a reference panel. We used the 1000 genome individuals with ancestry of European (N=379) released in December 2013 as the reference panel, imposed an LD threshold of 0.9 and selected a default setting based on a chi-square 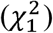 test statistic threshold of 10 for SNP pruning. The GWAS summary data were based on the meta-analysis results at the combined stage. As a sensitivity check, we repeated the HEIDI analysis using the summary GWAS data of individuals with European ancestry (studies 1-11).

### Telomere length association studies for Mendelian randomization analysis

We gathered the summary statistics from three large-scale meta-analysis studies for association with LTL, including (I) the association results of 484 SNPs located in *TERT* locus listed in Supplementary Info from the study conducted by Bojesen et al.^24^ (N=53,724 individuals of European ancestry), (II) the GWAS summary data from the study conducted by Codd et al.^25^ (N=37,684 individuals of European ancestry) downloaded from the European Network for Genetic and Genomic Epidemiology consortium (ENGAGE, see URL) and (III) the association results of 4 SNPs at P < 5.0×10^-8^ and 1 SNPs at P < 1.0×10^-6^ listed in Table 1 from the study conducted by Pooley et al.^26^ (N=26,089 individuals of European ancestry). In the first study, the effect sizes were available for change in telomere length (Δ*TL* > 0 indicating a test allele associated with longer LTL) and fold change in telomere length (>1 indicating a test allele associated with longer LTL). We used the effect sizes with respect to Δ*TL* in our analysis. The other two studies reported Δ*TL* in their summary data. All telomere lengths refer to the relative telomere to single copy gene (T/S) ratios using quantitative PCR methods with different scaling approaches applied to each study. The summary data of the second were used for bi-directional Mendelian randomization analysis. Summary statistics of IEAA and EEAA were based on the association results from the combined meta-analysis. Our SMR analysis used the summary data from studies I & II. To compare the patterns between LTL associations and IEAA associations at 5p15.33 *TERT* locus, we used the dense panel of SNP association results from study I (484 SNPs), as depicted in **Figure 3b**.

### IEAA association in brain tissues

To study the generalization of the overlap between telomere association and IEAA association in non-blood tissues, we used the GWAS results of IEAA in brain tissues from our recent studies including brain regions in cerebellum (N=555)^18^, in prefrontal cortex (N=657)^70^ and in multiple areas across cerebellum, frontal cortex, pons, temporal cortex and prefrontal cortex (N=1,796 brain tissues)^70^. All the study populations were of European ancestry. All the IEAA measures were derived from Horvath DNAm Age. But, strictly speaking, the measure variable of age acceleration in cerebellum was simply based on the residuals from regression DNAm Age on chronological age only, referred as to *AgeAccelerationResidual* which is subtly distinct from IEAA.

### In vitro studies of hTERT in human fibroblasts

Foreskins were obtained from routine elective circumcision. The tissue was cut into small pieces and digested overnight at 4C with Liberase, after which the epidermis was peeled away from the dermis. Several of dermis were placed face down in a plastic dish with DMEM supplemented with 10% foetal calf serum, penicillin, streptomycin and gentamycin. After incubation at 37oC with 5% carbon dioxide for a week, fibroblasts that emerged from the tissues were harvested and expanded in fresh vessels.

Recombinant retroviruses bearing the hTERT gene (pBabePurohTERT) or empty vector (pBabePuro) were prepared by transfecting Phoenix A cells, recovering the recombinant viruses in the media and using them to infect primary fibroblasts. Following selection with 1ug/ml puromycin, surviving cells were used in the experiment described.

### GWAS based enrichment analysis with MAGENTA

We used the MAGENTA software ^28^ to assess whether our meta-analysis GWAS results of epigenetic age acceleration are enriched for various gene sets, e.g. KEGG pathways, Gene Ontology (GO) terms such as biological processes or molecular functions. To assign genes to SNPs, we extended gene boundaries to +/- 50kb. For computational reasons, we removed categories that did not contain any genes related to age acceleration at a level of P<1.0×10^-3^ or that contained fewer than 10 genes. The cutoffs of gene set enrichment analysis (GSEA) in the MAGENTA algorithm were set at 95^th^ and 75^th^ percentiles which are the default parameter values for a general phenotype and for a highly polygenic trait, respectively ^28^.

Initially, empirical *P* values were estimated based on 10,000 permutations. For significant gene sets (empirical *P* < 1.0×10^-4^), we estimated the final empirical P value using one million permutations. We only report gene sets whose false discovery rate FDR (calculated by MAGENTA) was less than 0.20.

### LDSC genetic correlation analysis

We performed cross-trait LD score regression^29^ to relate IEAA/EEAA to various complex traits (27 GWAS summary data across 23 distinct phenotypes). The GWAS results for IEAA and EEAA were based on the summary data at *stage 1* analysis.

The following is a terse description of the 27 published GWAS studies. Two GWAS results in individuals of European ancestry came from the GIANT consortium on body fat distribution: waist circumference and hip to waist ratio. GWAS results of BMI and height also from the GIANT consortium. Further, we used published GWAS results from inflammatory bowel disorder (IBD) and its two subtypes: Crohn’s disease and ulcerative colitis, lipid levels, metabolic outcomes and diseases: insulin and glucose levels, type 2 diabetes (stage 1 results) phenotype, age-related macular degeneration (neovascular and geographic atrophy), Alzheimer’s disease (stage 1 results), attention deficit hyperactivity disorder (ADHD), bipolar disorder, major depressive disorder, schizophrenia, education attainment, age at menarche, age at menopause, LTL and longevity. The summary data of LTL was based the GWAS conducted by Codd et al^25^, as described in an earlier section. A description of other published GWAS study can be found in **Supplementary Note 4**.

As recommend by LDSC, we filtered to HapMap3 SNPs for each GWAS summary data, which could help align allele codes of our GWAS results with other GWAS results for the genetic correlation analysis conducted. We constrained intercepts with the --intercept –h2 flag for the GWAS studies with GC-correction that we input the estimates of intercepts obtained from the heritability analysis implemented under LDSC.

### MR-Egger regression

Under a weaker set of assumptions than typically used in Mendelian randomization (MR), an adaption of Egger regression can be used to detect and correct for the bias due to directional pleiotropy ^27^. While the standard method of MR estimation, two-stage least squares, may be biased when directional pleiotropy is present, MR-Egger regression can provide a consistent estimate of the causal effect of an exposure (e.g. age at menopause) on an outcome trait (e.g. epigenetic age acceleration). In testing the regression model, we used the leading variants (P < 5.0×10^-8^ or their surrogates) from each GWAS locus associated with the exposure, as instrumental variables. We performed LD-based clumping procedure in PLINK with a threshold of r^2^ set at 0.1 in a window size of 250kb to yield the leading variants present in both GWAS summary data sets (for exposure and outcome), as needed. The random effects model meta-analysis was performed using “MendelianRandomization” R package.

### GWAS-based overlap analysis between age acceleration and various phenotypes

Our GWAS-based overlap analysis related gene sets found by our GWAS of epigenetic age acceleration with analogous gene sets found by published GWAS of various phenotypes. We used the MAGENTA software to calculate an overall GWAS *P* value per gene, which is based on the most significant SNP association *P* value within the gene boundary (+/- 50 kb) adjusted for gene size, number of SNPs per kb, and other potential confounders ^28^. To assess the overlap between age acceleration and a test trait, we selected the top 2.5% (roughly 500 genes ranked by P values) and top 10 % genes (roughly 1900 genes) for each trait and calculated one-sided hypergeometric P values ^18,70^. In contrast with the genetic correlation analysis, GWAS based overlap analysis does not keep track of the signs of SNP allele associations.

We performed the overlap analysis for all the 23 complex traits used in the genetic correlation LDSC analysis and a few more studies that we were not able to conduct the LDSC analysis due to small sample size (N < 5000), negative heritability estimates or the entire study population from non-European ancestry. The additional traits including modifiers of Huntington’s disease motor onset, Parkinson’s disease and cognitive functioning traits (**Supplementary note 4**).

## URLs

1000 genome project, http://www.1000genomes.org/

BRAINEAC, http://www.braineac.org/

DNAm age, <u>http://labs.genetics.ucla.edu/horvath/htdocs/dnamage/</u>

ENGAGE, https://downloads.lcbru.le.ac.uk/

EIGENSTRAT, http://genepath.med.harvard.edu/~reich/Software.htm

GTEx, http://www.gtexportal.org/home/documentationPage#AboutGTEx

GIANT, https://www.broadinstitute.org/collaboration/giant/index.php/Main_Page

HaploReg, http://www.broadinstitute.org/mammals/haploreg/haploreg.php

HRC, http://www.sanger.ac.uk/science/collaboration/haplotype-reference-consortium

HRS, http://hrsonline.isr.umich.edu/

Illumina Infinium 450K array annotation, http://support.illumina.com/downloads/infinium_humanmethylation450_product_files.html. IMPUTE2, https://mathgen.stats.ox.ac.uk/impute/impute_v2.html

LD score Regression, https://github.com/bulik/ldsc

Locuszoom, http://csg.sph.umich.edu/locuszoom/

METAL, http://csg.sph.umich.edu/abecasis/Metal/

MAGENTA, https://www.broadinstitute.org/mpg/magenta/

NESDA, http://www.nesda.nl/en/

NTR, http://www.tweelingenregister.org/en/

PLINK, http://pngu.mgh.harvard.edu/~purcell/plink/

R metafor, http://cran.r-project.org/web/packages/metafor/

R WGCNA, http://labs.genetics.ucla.edu/horvath/CoexpressionNetwork/

SHAPEIT, https://mathgen.stats.ox.ac.uk/genetics_software/shapeit/shapeit.html

SNPTEST, https://mathgen.stats.ox.ac.uk/genetics_software/snptest/snptest.html

YFS, http://youngfinnsstudy.utu.fi/index.html

## ACKNOWLEDGEMENTS

The study was supported by U34AG051425-01 and NIA/NIH 5R01AG042511-02. The WHI program is funded by NIH/NHLBI, U.S. Department of Health and Human Services through contracts NIH/NHLBI 60442456 BAA23 (Assimes, Absher, Horvath), HHSN268201100046C,HHSN268201100001C,HHSN268201100002C, HHSN268201100003C, HHSN268201100004C, and HHSN271201100004C, Epigenetic Mechanisms of PM-Mediated CVD Risk (WHI-EMPC), NIH/NIEHS R01ES020836 (Whitsel, Baccarelli; Hou). SNP Health Association Resource project (WHI-SHARe), NIH/NHLBI N02HL64278 GWAS of Hormone Treatment and CVD and Metabolic Outcomes within the Genomics and Randomized Trials Network” (WHI-GARNET), NIH/NHGRI U01HG005152 (Reiner). The Framingham Heart Study is funded by National Institutes of Health contract to Boston University N01-HC-5195 and HHSN268201500001I and its contract with Affymetrix, Inc for genotyping services (Contract No. N02-HL-6-4278). The DNA methylation resource from the FHS was funded by the Division of Intramural Research, National Heart, Lung, and Blood Institute, National Institutes of Health. LX, KLL and JMM were supported by R01AG029451. DPK was supported by R01AR041398 and R01AR061162. TwinsUK is funded by the Wellcome Trust and European Community’s Seventh Framework Programme (FP7/2007-2013), and also receives support from the National Institute for Health Research (NIHR)-funded BioResource, Clinical Research Facility and Biomedical Research Centre based at Guy’s and St Thomas’ NHS Foundation Trust in partnership with King’s College London. SNP Genotyping in TwinsUK was performed by The Wellcome Trust Sanger Institute and National Eye Institute via NIH/CIDR. PCT and JTB were supported by ESRC (ES/N000404/1).

## CONTRIBUTIONS

ATL carried out all of the non-study-specific statistical analyses and wrote the first draft of the article. LX performed most FHS specific analyses and preliminary meta-analyses. All authors participated in the interpretation of the study and helped revise the article. SH conceived of the study.

## COMPETING FINANCIAL INTERESTS

The authors declare no competing financial interests.

## CORRESPONDING AUTHOR

Correspondence to Steve Horvath

